# Gender imbalances in citation rates are mediated by field specific author gender distributions

**DOI:** 10.1101/2022.07.28.501862

**Authors:** Steffen Riemann, Mandy Roheger, Jan Kohlschmidt, Jennifer Kirschke, Margherita Lillo, Agnes Flöel, Marcus Meinzer

**Affiliations:** Department of Neurology, University Medicine Greifswald, Greifswald, Germany

**Keywords:** Gender bias, citation bias, speech and language pathology

## Abstract

Gender biases are well documented in science and typically favor male scientists. In this context, a particularly pervasive gender bias is undercitation of publications authored by women, resulting in profound negative effects on academic visibility and career advancement. This bias has been well documented in fields where author gender distributions are strongly skewed towards men (e.g., astronomy, physics, neuroscience). By investigating citation practices in a field that has traditionally been more accessible to female scientists (Speech and Language Pathology, SLP), we demonstrate that gendered citation practices are mediated by author gender distribution, rather than being a universal pattern. Specifically, our results revealed a citation pattern in SLP that (a) overall tends to favor female authors, (b) persists after controlling for potential confounding factors and, (c) is particularly strong when female authors are citing publications involving female first and senior author teams. This research also implies that the implementation of effective measures to increase the number and influence of underrepresented individuals in specific fields of science may be suitable to mitigate downstream disadvantages for career advancement of either sex.

## Introduction

Gender biases are well documented in academia and typically favor male researchers (Huang et al. 2020; Ross et al. 2022). For example, female compared to male researchers frequently fare worse regarding work recognition and compensation, grant funding outcomes, teaching evaluations, hiring, or tenure promotion. Moreover, despite increasing awareness of gender biases in academia and initiatives to address them, necessary change appears to be slow and substantial inequality remains (Llorens et al. 2021).

In this context, undercitation of female scholars is a particularly detrimental type of gender bias that has profound negative effects on academic visibility and career advancement (Dworkin et al. 2020; Llorens et al. 2021). Several recent studies have demonstrated that reference lists of scientific publications tend to include more papers with men as first and last authors than one would expect based on the gender distribution of authors in specific fields. This pattern has consistently been shown across several fields, including neuroscience (Dworkin et al. 2020; Fulvio et al. 2021), physics (Teich et al. 2021), astronomy (Caplar et al. 2017), international relations (Maliniak et al. 2013), and political science (Sa et al. 2020). These studies suggest that male overcitation is a pervasive and universal pattern in science, with widespread negative implications for female scholars.

However, previous studies on citation bias in science have exclusively investigated fields with author distributions strongly skewed towards male researchers. In these fields, sociological theory (McPherson et al. 2001) predicts that a combination of implicit or explicit favoritisms towards similar group members (e.g., individuals of the same biological sex, particularly by individuals in influential positions) and systemic factors (Dworkin et al. 2020; Llorens et al. 2021; Ross et al. 2022), contribute to overcitation of male scientists.

Therefore, the present study investigated citation bias for the first time in a field which has traditionally been more accessible to female scientists, i.e., speech-and language pathology (SLP). For example, males make up only 2.5% of speech- and language pathologists in the United Kingdom (Litosseliti & Leadbetter, 2013) and a recent survey in 13 countries revealed that on average, only 5% of registered speech and language therapists are male (range: 2-13%, retrieved 28/7/2022 from *https://twitter.com/speechguys*). Similarly, the percentage of male academics registered with different professional or scientific SLP organizations in 2021 ranged from 4.5-22%^1^. This field-specific pattern of gender distribution, and the resulting gender author distribution in SLP publications (see below), allows investigating if (male) citation bias is a universal phenomenon, even in fields that are less “dominated” by males, or is mediated by the author gender distribution in different fields.

## Results

The data for this study were drawn from the Web of Science (WoS) category “Audiology and Speech- and Language Therapy” and was based on 14 SLP journals (see methods). All analyses were conducted using an open-source code (Dworkin et al. 2020), adapted to the specific requirements of our study. To analyze historical changes of the author gender distribution in SLP publications, data from 1958-2021 were included. As in previous studies that used the same methodology (Dworkin et al. 2020; Fulvio et al. 2021, Teich et al. 2021, Wang et al. 2021), citation pattern-based analyses focused on the last decade (2010-2020; data from 2021 were not considered because of the small number of citations to these very recent articles). A total of 8.607 publications and 11.329 citations were included in the respective analyses.

### Trends in authorship

To investigate overall trends in authorship, we started with assigning gender categories (Men [M], Women [W]) to first and last authors’ first names using a probabilistic approach (see Methods). This creates four main categories for our analyses: man as first author, man as last author [MM]; man first, women last [MW]; women first, man last [WM]; women first, women last [WW]. In a first step, we plotted the four categories to visualize the categorical gender breakdown of authorship in the investigated journals from 1958 to 2021 (**Figure 1**), showing that the vast majority of publications in SLP in 1958 (69.2%) were authored by MM author teams. Over time, this proportion gradually decreased to 23.4% in 2021. Publications with at least one female author increased over time and in 2021, 21.2% of the publications were WM-authored, 19.1% were MW-authored. WW-authored publications increased to 36.3%. Hence, 76.6% of publications had at least one female author. For a detailed overview of the categorical gender breakdown in individual journals and available time frames see **Supplementary Figure 1**.

**Figure 1:**
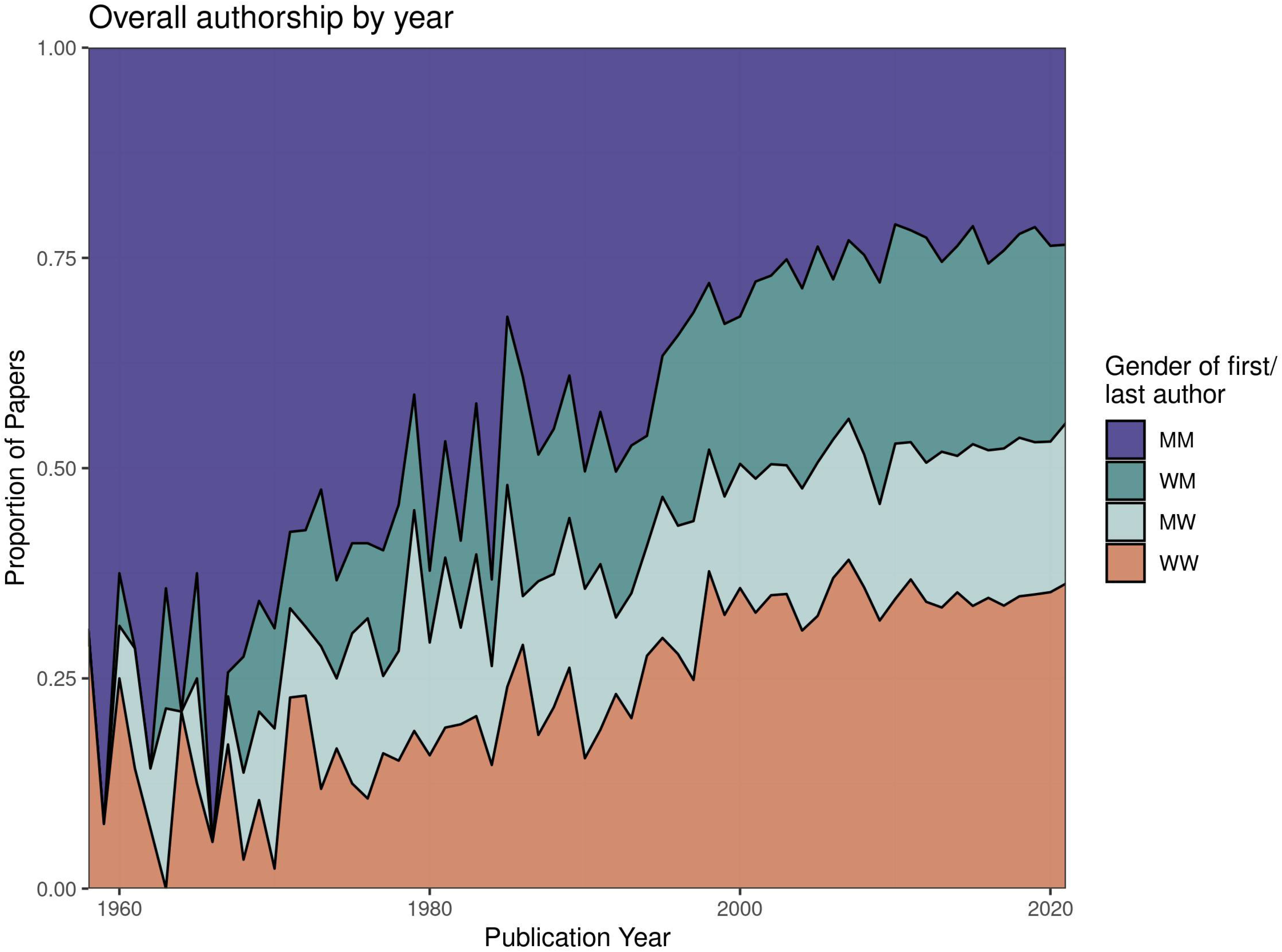
Gender breakdown averaged across all investigated journals and available time periods (1958 to 2021). Gender proportions are displayed in four categories: men as first and last authors (MM), women as first author and men as last author (WM), men as first author and women as last author (MW), and women as both first and last authors (WW)

### Citation analysis

To analyze citation behavior, we used the references lists of all manuscripts in the investigated journals between 2010 and 2020 using the four author categories (MM, MW, WM, WW). Two **Gender Citation Balance Indices** were calculated: (1) Gender Citation Balance Indices relative to all literature and (2) Gender Citation Balance Indices relative to the conditional citation gap (i.e., controlling for several potential confounding factors, see below). The time frame and measures were chosen to allow for comparison of our results to those recently reported in different fields of science (Dworkin et al. 2020; Fulvio et al. 2021, Teich et al. 2021, Wang et al. 2021).

(1) **Gender Citation Balance Indices relative to all literature** [e.g., the (percentage of MM citations observed - percentage of MM citations expected)/percentage of MM citations expected)] were calculated to compare author gender proportions in the reference lists to the overall gender proportion in the existing literature (see **Figure 2A** that illustrates over- and undercitation of different author gender groups, compared to their expected proportions using a random-draws model, Dworkin et al. 2020). Using this measure, MM publications were cited 21.2% less than expected (95%CI = (−22.5%, −19.8%)), WW publications were cited 13.3% more than expected (95%CI = (11.7%, 14.8%)). WM publications were cited 5.4% more than expected (95%CI = (3.8%, 6.9%)), MW publications were cited 8.6% more than expected (95%CI = (7.0%, 10.6%)).

**Figure 2:**
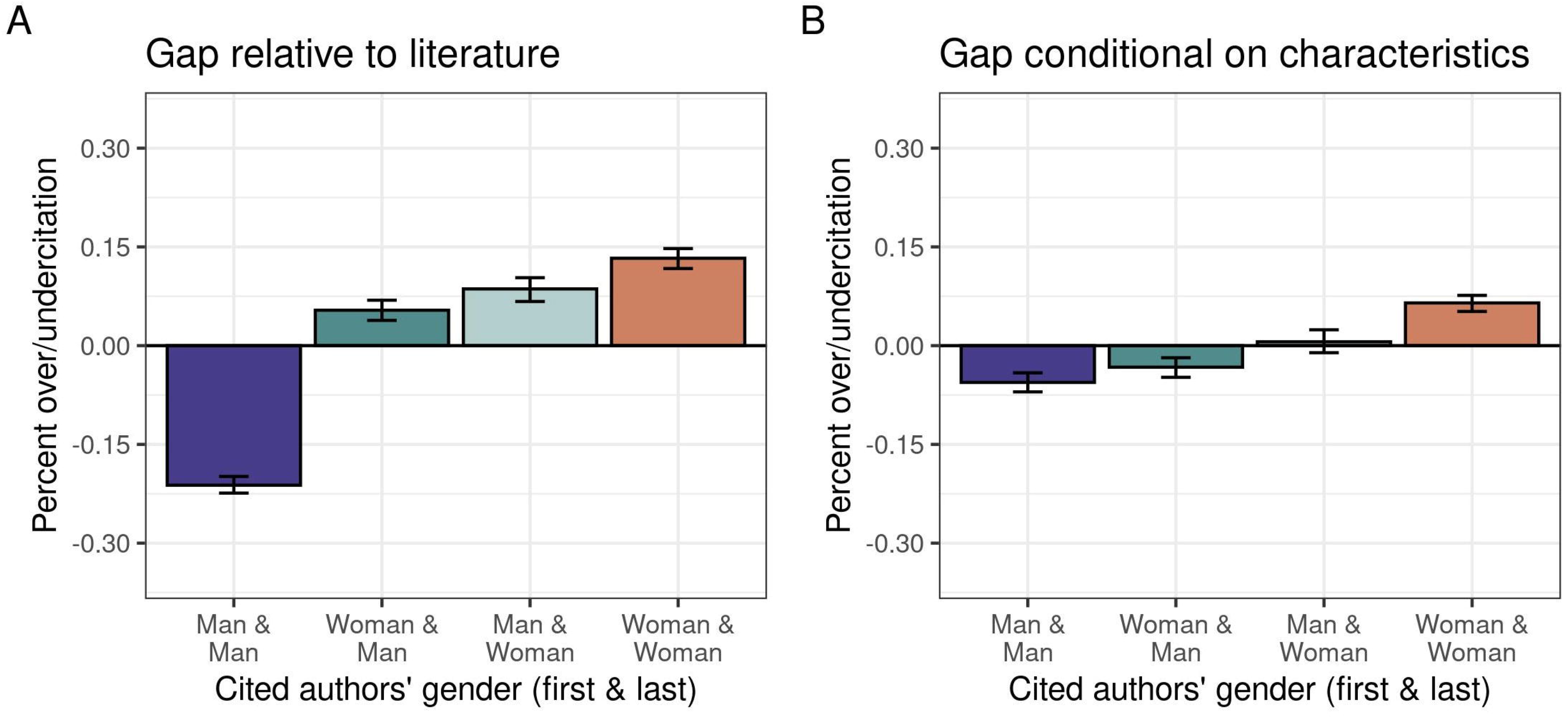
Measures of over- and undercitation. A. Illustrates the Gender Citation Balance Indices relative to all literature; B. Citation imbalance accounting for potentially relevant characteristics of publications (Gender Citation Balance Indices relative to the conditional gap). Positive and negative values represent over- and undercitation (in percent) for the different author categories (MM, WM, MW, WW)

(2) Comparison of citation rates to overall authorship proportions does not account for other factors that may affect citation behavior. Therefore, we calculated **Gender Citation Balance Indices relative to the conditional citation gap** using the same formula as above, but “expected percentages” were based on potentially relevant external factors (e.g., publication date, author count, seniority of first and last authors) using a generalized additive model (GAM, see Methods; detailed descriptions have been provided by Dworkin et al. 2020; Teich et al. 2021). This allows to calculate proportions of citations to the four author categories that would be expected if references were drawn in a genderblind manner from pools of publications sharing similar characteristics. Using this approach, MM publications were cited 5.6% less than expected (95%CI = (−7.1%, −4.2%)), WW publications were cited 6.5% more than expected (95%CI = (5.2%, 7.6%)). WM publications were cited 3.2% less than expected (95%CI = (−4.6%, −1.9%)), and MW publications were cited 0.5% more than expected (95%CI = (−1.2%, 2.3%); **Figure 2B)**.

(3) In a final analysis, we investigated the **effect of author gender on citation behavior** (see **Figure 3**). Therefore, we used the Gender Citation Balance Indices relative to the conditional gap, and plotted them separately by the gender of the citing authors. This revealed that within MM reference lists (**Figure 3B**), MM publications received more citations (2.7%, 95%CI = −0.5%, 0.6%) than expected, MW publications were cited 1.5% more than expected (95%CI = −0.5%, 0.1%; **Figure 3D**), WW publications were cited 1.3% (95%CI = −0.4%, 0.1%) less than expected (**Figure 3E**), and WM publications were cited 2.6% less than expected (95%CI = −7.3%, 2.7%; **Figure 3C)**. A different pattern occurred within WW reference lists, in which other WW publications were cited 10.4% (95%CI = 0.8%, 12.5%) more than expected and MM publications were cited 9.7% less than expected (95%CI = −11.8%, −7.1%). WM publications were cited 4.1% (95%CI = −6.3%, −1.7%) less than expected, MW publications were cited 1.2% (95%CI = −3.9%, 1.9%) less than expected. Within WM reference lists, MM publications were cited 6.9% (95%CI = −4.0%, 9.7%) less than expected, WM publications were cited 4.8% (95%CI = −7.7%, −2.0%) less than expected, MW publications were cited 1.5% (95%CI = −1.9%, 5.4%) more than expected, and WW publications were cited 8.3% (95%CI = 5.6%, 11.0%) more than expected. Within MW reference lists, MM publications were cited 6.4% (95%CI = −9.8%, −2.9%) less than expected, WM and MW publications were cited 0.2% (95%CI = −3.7%, 3.2%) less and 1% (95%CI = −2.9%, 5.1%) more than expected and WW publications were cited 4.7% (95%CI = 1.7%, 7.9%) more than expected.

**Figure 3:**
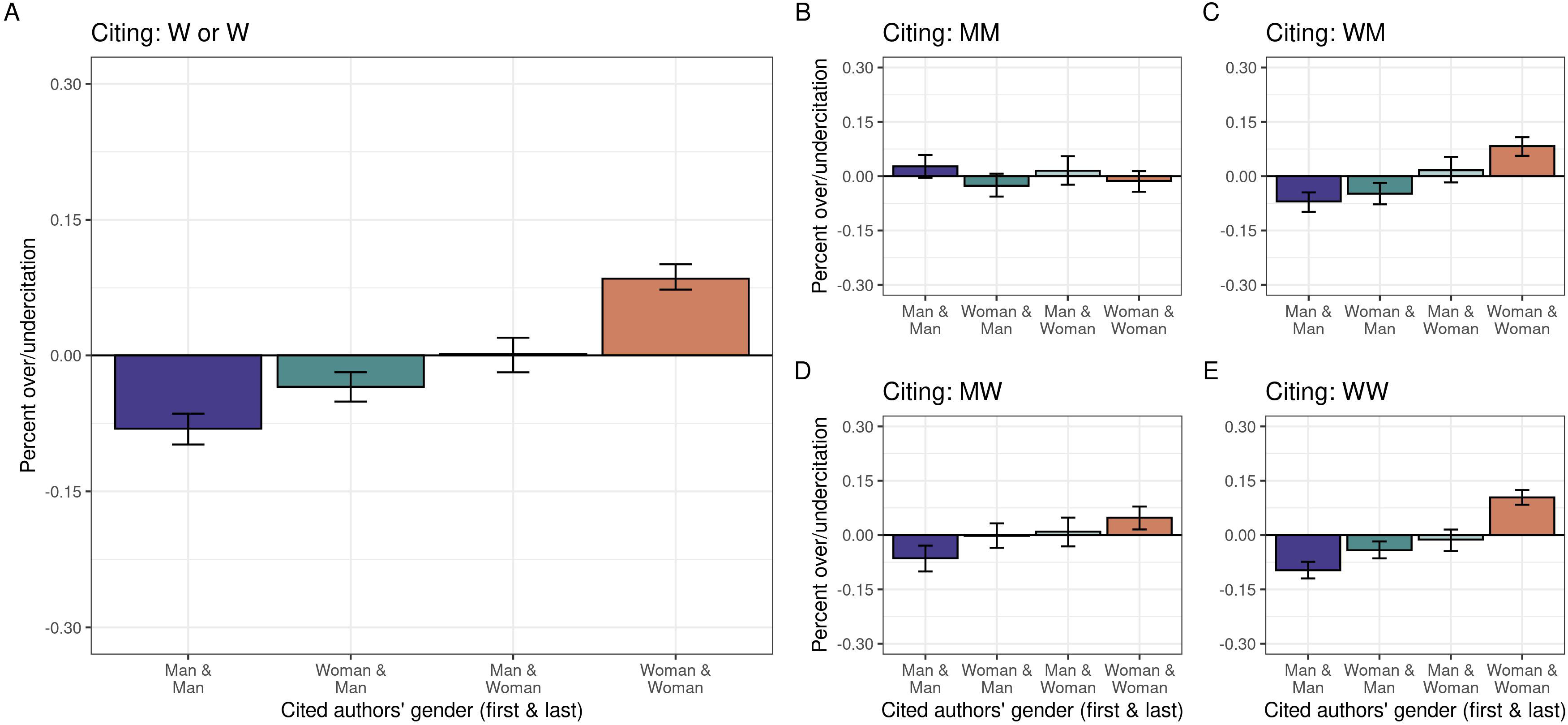
Relationship between author gender and gendered citation practices. A. Illustrates Gender Citation Balances relative to the conditional gap for citations involving at least one female author (W or W). B-E. Show citing behavior of authors from the four author categories (MM, WM, MW, WW)

None of these percentages of over- and undercitation are statistically significant after correcting for multiple comparisons. However, when aggregating all publications, in which women are involved as authors, either as first, last or both, publications with female first and last authors were significantly overcited by 8.5% (95%CI = 7.3%, 10.1%; *p* = 0.015; **Figure 3A)**. MM author teams were undercited by 8.1%, yet this result did not reach significance (95%CI =-9.8%, −6.4%, *p* = 0.13).

## Discussion

Our results show that the author gender distribution in SLP publications is currently more balanced compared to previously investigated fields of science that are largely dominated by male authors (Dworkin et al. 2020; Fulvio et al. 2021, Teich et al. 2021, Wang et al. 2021). Moreover, for the last decade, our results demonstrate a citation pattern in SLP that (a) overall tends to favor female authors, (b) persists after controlling for potential confounding factors and, (c) is particularly strong for publications involving female first and last author teams. While male citation bias appears to be common in many fields of science (Llorens et al. 2021), our results challenge the notion that this is a universal pattern and rather emphasize the contribution of author gender distributions in specific sub-fields of science.

Previous studies interrogating citation bias have highlighted a pervasive pattern of undercitation of female scholars (Maliniak et al. 2013; Caplar et al. 2017; Dworkin et al. 2020; Sa et al. 2020; Fulvio et al. 2021; Teich et al. 2021, Wang et al. 2021). However, these studies have exclusively investigated fields where author distributions are strongly skewed towards men (e.g., neuroscience, physics or astronomy). Even in recent years, the majority of publications in these fields were mainly contributed by male authors. The most dramatic example comes from contemporary physics, where only 3.6% of articles were contributed by WW author teams and only 33% involved at least one female first or senior author (Teich et al. 2021). Indeed, similar author team distributions have been reported in all previously investigated fields to date. For example, in neuroscience, 78-90% of publications involved male authors, either as first or senior author, or both (Dworkin et al. 2020; Fulvio et al. 2021). In contrast, the present study investigated for the first time citation patterns in a field where the majority of publications with prominent authorship positions are currently contributed by women (i.e., ~75% of publications with a woman as first and/or senior author over the last two decades, including ~30% WW author teams, see **Figure 1**). For this specific field, we demonstrate that author teams involving at least one woman are overciting publications authored by female first and last author teams (WW) by almost 10%. This suggests that the degree of undercitation of women (and potential negative downstream effects on visibility and career advancement) is at least partly rooted in authorship distributions of specific fields of science, rather than being a universal pattern.

Two main limitations of this research need to be acknowledged: As in previous studies (Dworkin et al. 2020; Fulvio et al. 2021, Teich et al. 2021, Wang et al. 2021), first names of authors were identified using a probabilistic and binary gender algorithm, which may not reflect the actual self-identification of authors. However, because we used the same algorithm as in previous publications, the overall results of this study are still comparable to previous work. Moreover, self-identification as male, female or diverse is rather negligible in the present study, because the relevant factor is the gender identity assigned by the citing authors. Another limitation is the relatively small field that was investigated (SLP), which resulted in limited statistical power (i.e., only aggregated effects reached statistical significance) and also prevented investigation of temporal trends in authorship across the past decade. Nonetheless, the overall pattern of the results in a field that is largely dominated by female scientists, are consistent with those reported previously in fields dominated by men (Dworkin et al. 2020; Fulvio et al. 2021, Teich et al. 2021, Wang et al. 2021), even though the direction of the effects (favoring female vs. male scientists, respectively) was different. This suggests that overcitation critically depends on authorship distribution in specific fields, which also has direct implications for potential interventions aimed at reducing gender based bias in science.

Specifically, while the exact mechanisms underlying (male or female) citation bias are currently unknown and the present study was not designed to pinpoint the many potential individual and systemic factors that mediate the observed undercitation of female or male authors (for discussions see Dworkin et al. 2020; Llorens et al. 2021), the present study highlights the importance of a common human factor to citation bias in science (i.e., “homophily”, McPherson et al. 2001). This concept describes the tendency of individuals to associate, bond or (implicitly or explicitly) favor other individuals that are similar with regard to age, race, and social status, but also gender. Within this framework, it would be expected that scientific fields dominated (either with regard to numbers or degree of influence on a given field) by men or women alike, show a bias towards citing work (or “appreciation”, Ross et al. 2022) of their own “gender in-group”. In the broader context of the current literature on disadvantages female scholars are faced with (Llorens et al. 2021), this theory provides not only an explanation for the currently reported patterns of citation bias, but also suggests that implementation of effective measures aimed at increasing the percentage of underrepresented individuals or connecting them to influential scholars (Verhoeven et al. 2020), may indeed be suited to leveling the playing field.

In sum, our results challenge the notion that male citation bias is a universal phenomenon in science and highlight the contribution of field specific author gender distribution to citation bias. Implementing effective measures to increase the number and influence of underrepresented individuals in specific fields of science may allow mitigation of downstream disadvantages regarding visibility and career advancement.

## Materials and Methods

### Data collection

Data analyses were based on the following journals included in the WoS category “Audiology and Speech- and Language Therapy”: American Journal of Speech and Language Pathology (available from 1998-2020), Aphasiology (1988-2020), Augmentative and Alternative Communication (2005-2020), Communication Sciences and Disorders (2015-2020), Folia Phoniatrica et Logopaedica (1994-2020), International Journal of Language & Communication Disorders (1998-2020), International Journal of Speech Language Pathology (2008-2020), Journal of Communication Disorders (1967-2020), Journal of Fluency Disorders (1977-2020), Journal of Speech Language and Hearing Research (1997-2020), Journal of Voice (1990-2020), Language and Speech (1958-2020), Language Speech and Hearing Services in Schools (1995-2020), and Seminars in Speech and Language (2012-2020). Journal selection was also guided by a short email survey asking 10 academics working in speech- and language therapy for journals representing their field. Journals from audiology and those publishing mainly experimental or neuroscientific research (e.g., Brain and Language) and general medical journals were not considered.

We downloaded all available published articles between 1958 and 2021 and included original articles, review articles and proceeding papers that were labeled with a digital object identifier (DOI). The data downloaded for each paper included author names, reference lists, publication dates and DOIs, and we obtained information referencing behavior by matching DOIs contained within a reference list to DOIs of papers included in the dataset. When no first names of authors could be obtained from the published papers, we searched them using CrossRef API (*www.crossref.org*).If first names were not available on CrossRef, we searched for them on the journal webpage. Dworkin et al. (2020) implemented a name disambiguation algorithm to minimize the number of papers for which we only had access to author’s initials, which was also used in the present paper (see Dworkin et al., 2020 for detailed information on the algorithm).

### Determination and interpretation of authors’ gender

If first names of authors were available or could be obtained using the algorithm, gender was assigned to first names using the “gender” package in R with the Social Security Administration baby name data set. For names that were not included in the R package, gender was assigned using *https://genderize.io/*, a freely available service that contains roughly 250.000 names. Please note that in the original study by Dworkin et al., (2020) a different, paid service was used (http://gender-api.com/). As suggested in the literature, we assigned ‘man’ (‘woman’) to each author if their name had a probability ≥0.70 of belonging to someone labeled as ‘man’ (‘woman’). Overall, gender could be assigned to both the first and last authors in 89.51% of the papers in our dataset. Of the 10.49% of papers with missing data, the first or last author’s name either had uncertain gender (6.67%) or was not available (3.82%). When interpreting our data please note, that the term “gender” does not directly refer to the sex of the author, as assigned by birth or chosen later, nor does it directly refer to the gender of the author, as socially assigned or self-chosen. It is important to keep in mind that the term “gender” in the present analysis is a function of the probability of assigned gendered names; the actual sex or gender of the authors is not and cannot be identified with this method.

### Statistical Analysis

For detailed information, code of the statistical analysis and explanatory formulas please refer to Dworkin et al. (2020). The original code that was adapted for the present study is available at *https://github.com/idwor/gendercitation*.

By assigning gender to authors’ first names of the manuscripts and citations, we created four main categories for our analyses: man as first author, man as last author [MM]; man first, women last [MW]; women first, man last [WM]; women first, women last [WW]. We then computed two Gender Citation Balance Indices for each of the four gender categories (i.e., WW, MM, WM, MW): (1) Gender Citation Balance Indices relative to all literature and (2) Gender Citation Balance Indices relative to the conditional citation gap. Gender Citation Balance Indices relative to all literature was computed as:

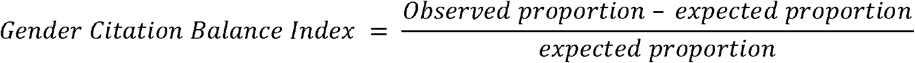

Positive values correspond to more frequent citations than expected and negative values correspond to less frequent citations than expected, respectively. Gender Citation Balance Indices relative to the conditional citation gap were computed in a similar manner, however, to obtain expected rates that account for various characteristics that may be associated with citation rates, we fit a generalized additive model on the multinomial outcome (MM, MW, WW, WM) using R package *“mgcv”*. Following the methodology reported by Dworkin et al. (2020), the model’s features were (1) year of publication, (2) the combined number of publications by the first and last authors (seniority), (3) the number of authors on the paper, (4) the journal in which it was published and (5) the type of publication (e.g., original article of review). When this model is then applied to each paper, it yields a set of probabilities that the paper belongs to the MM, WM, MW, and WW categories and citation rates can be predicted as if the citation was independent of the authors’ gender. Replicating the code provided by Dworkin et al. (2020) allows for direct comparison of our results to those of other studies in different fields of science that used the same approach (Dworkin et al. 2020; Fulvio et al. 2021, Teich et al. 2021, Wang et al. 2021).

## Acknowledgements

We would like to thank Jordan Dworkin for assistance with adapting the code that was used for the respective analyses.

## Competing Interests

The authors report competing interests.

## Footnotes

1 Personal communication with representatives of Speech Pathology Australia, American Speech, Language and Hearing Organization, The Academy of Aphasia and The German Association for Aphasia Research and Treatment.

## Notes

### Competing Interest Statement

The authors have declared no competing interest.

