## Supplementary Information 1 for "Gender imbalances in citation rates are mediated by field specific author gender distributions"

Categorical author gender breakdown and available time frames for the included journals (Prop = Proportion of male/female author teams in the the four categories, MM = man/man, WM = woman/man, MW = man/woman, WW = woman/woman)

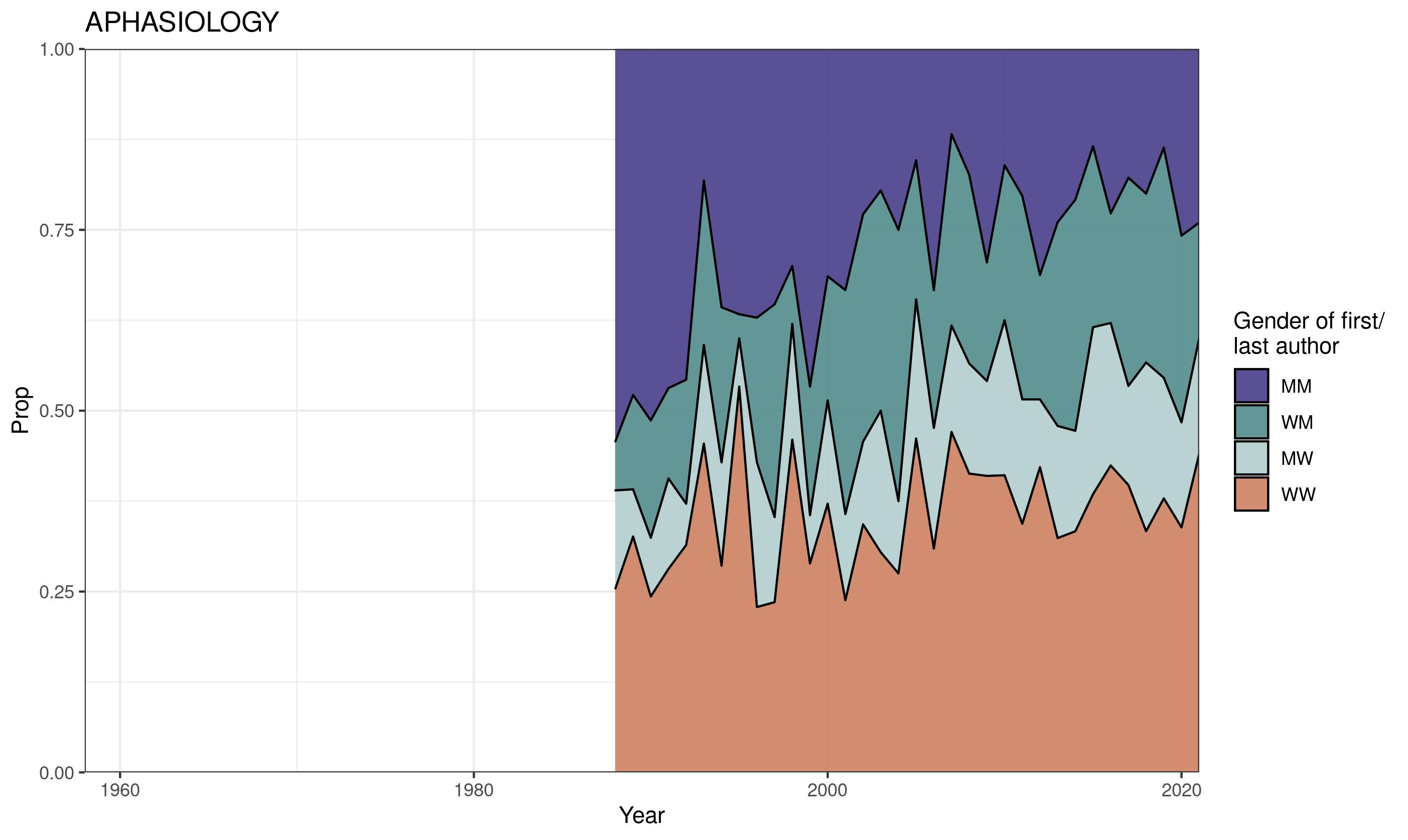

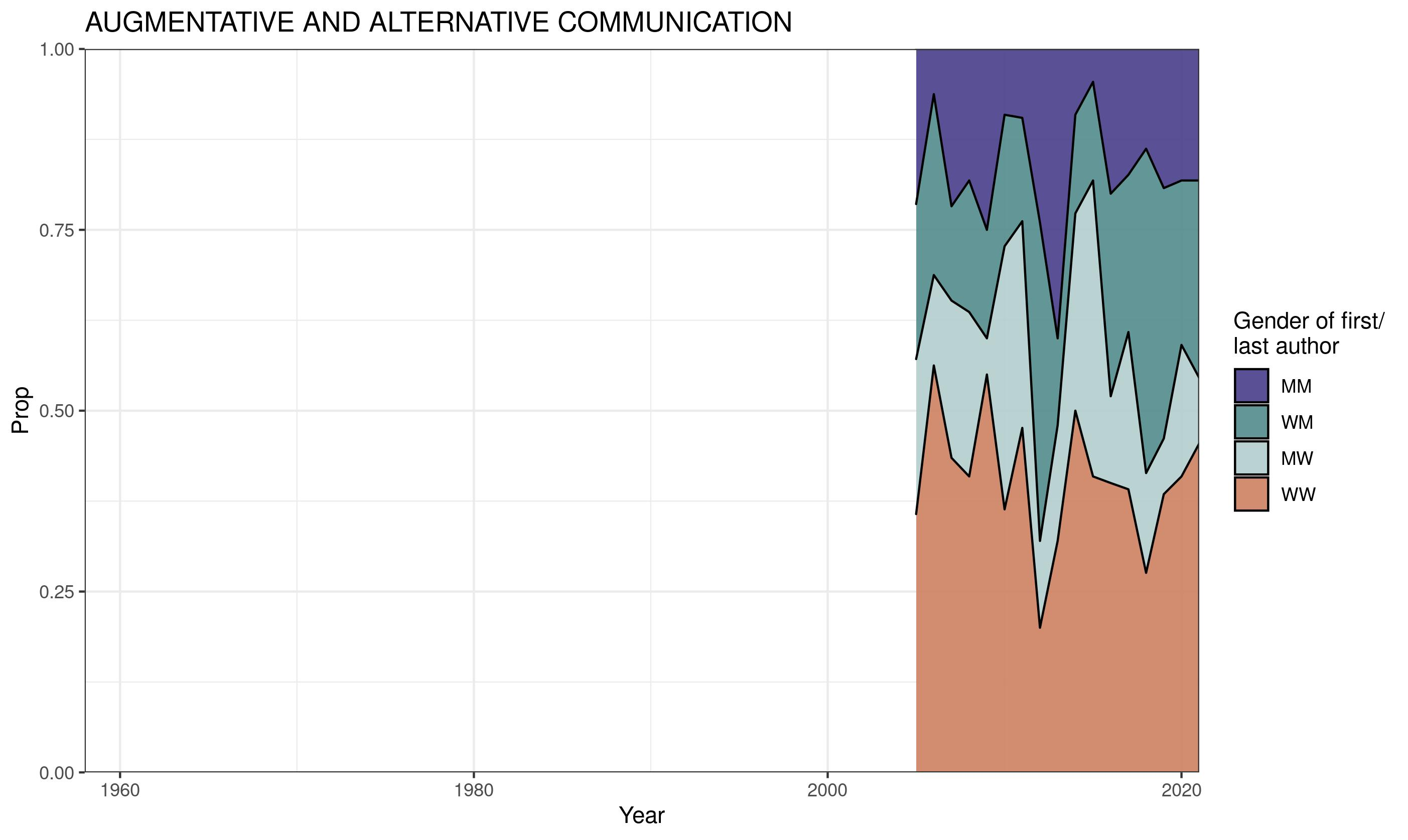

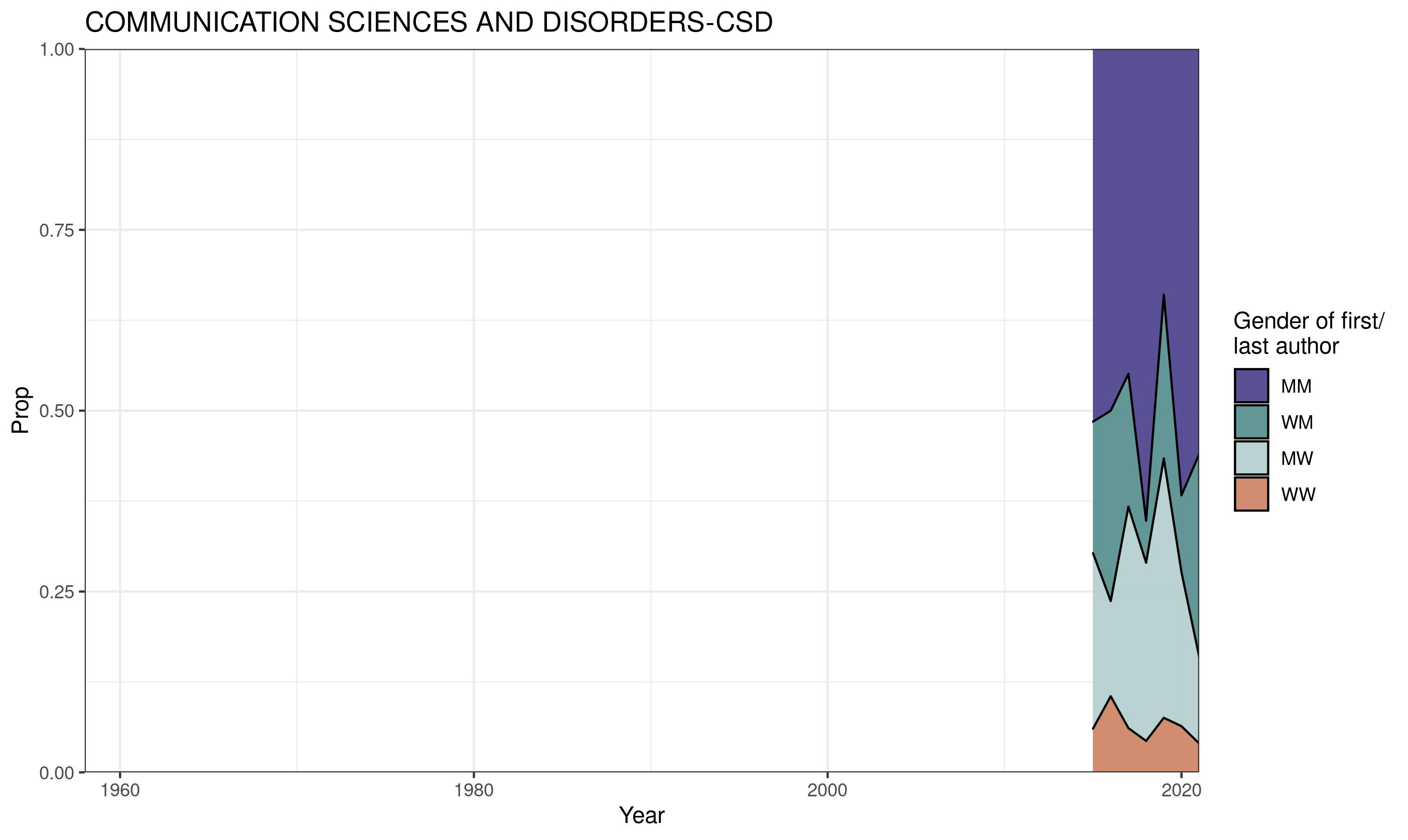

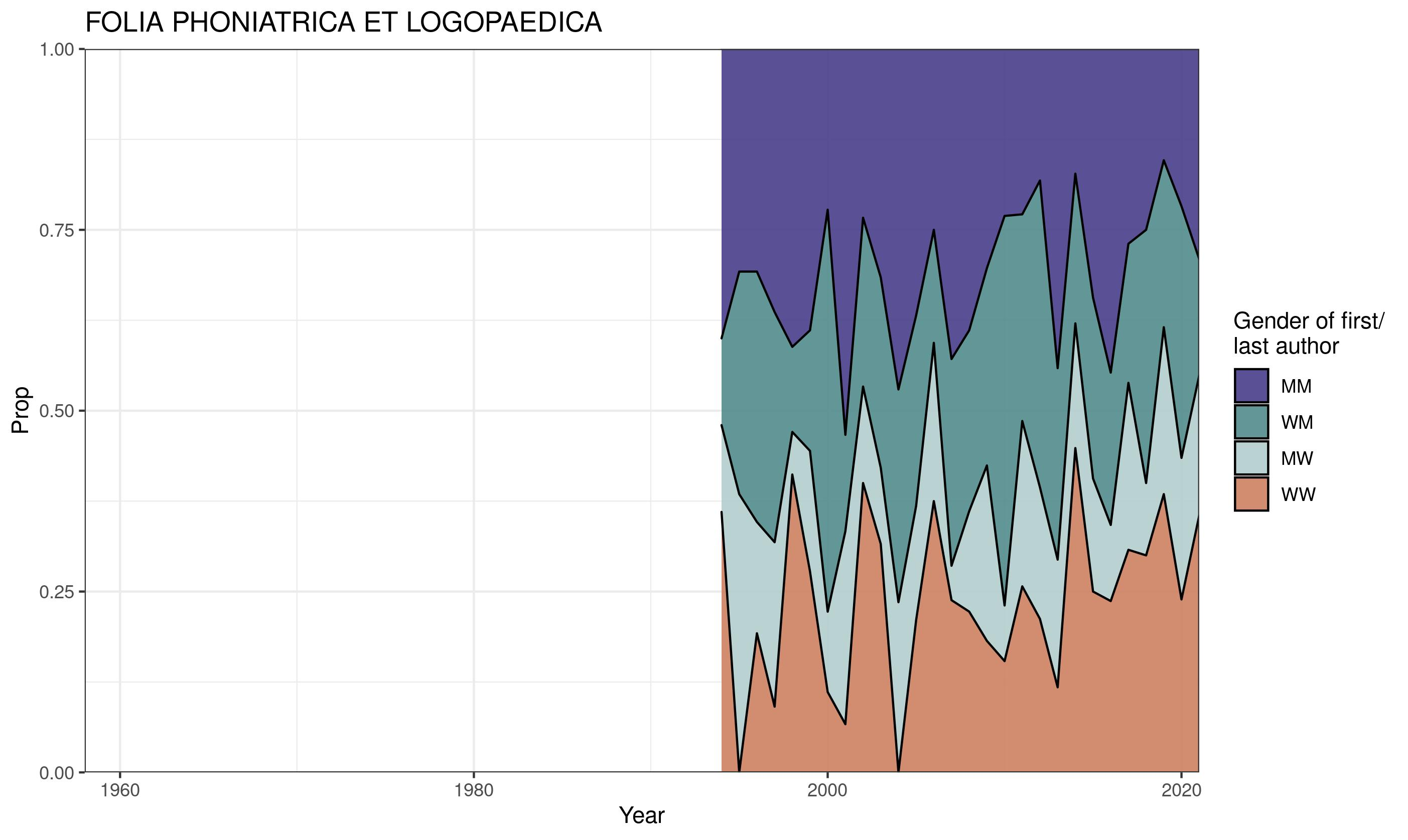

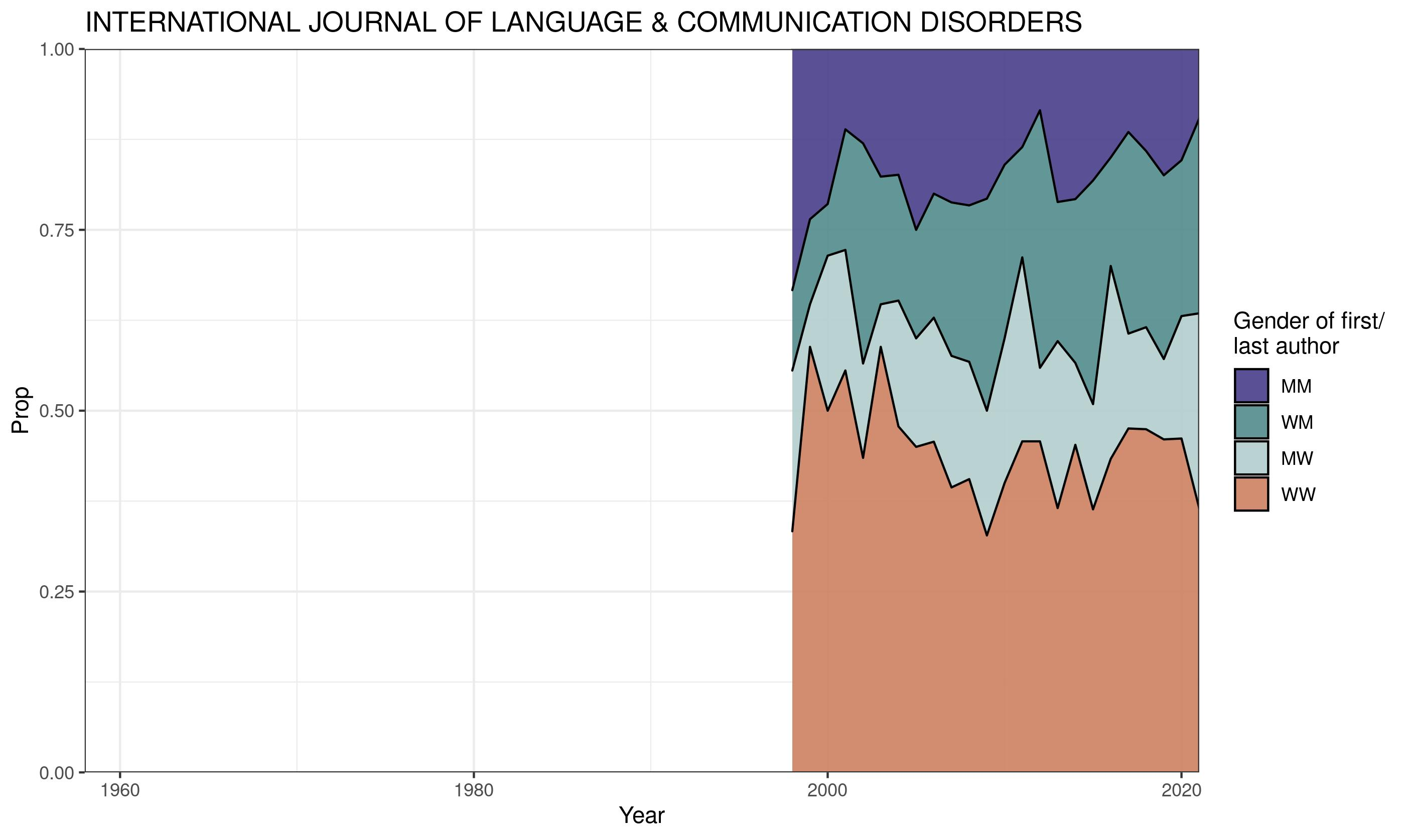

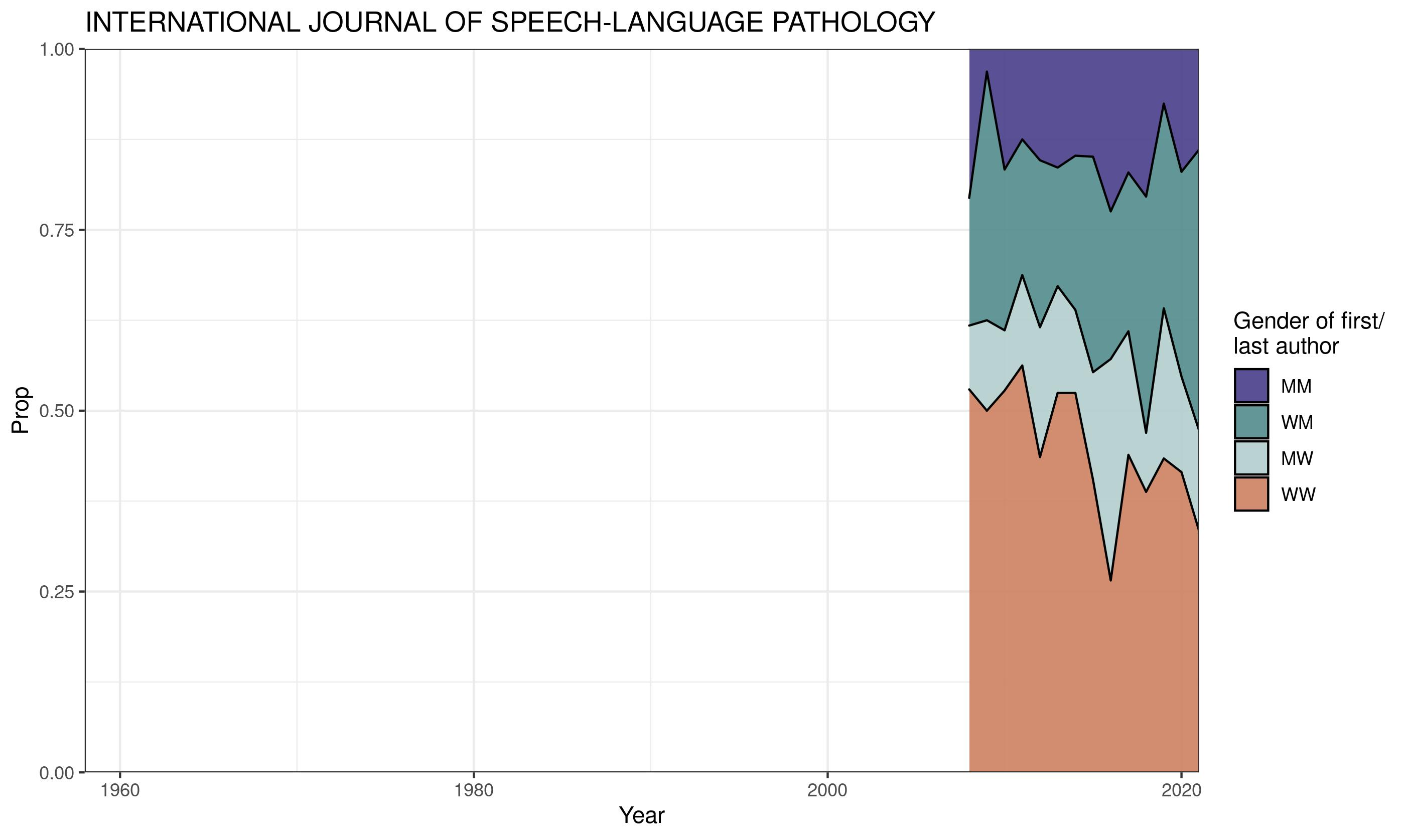

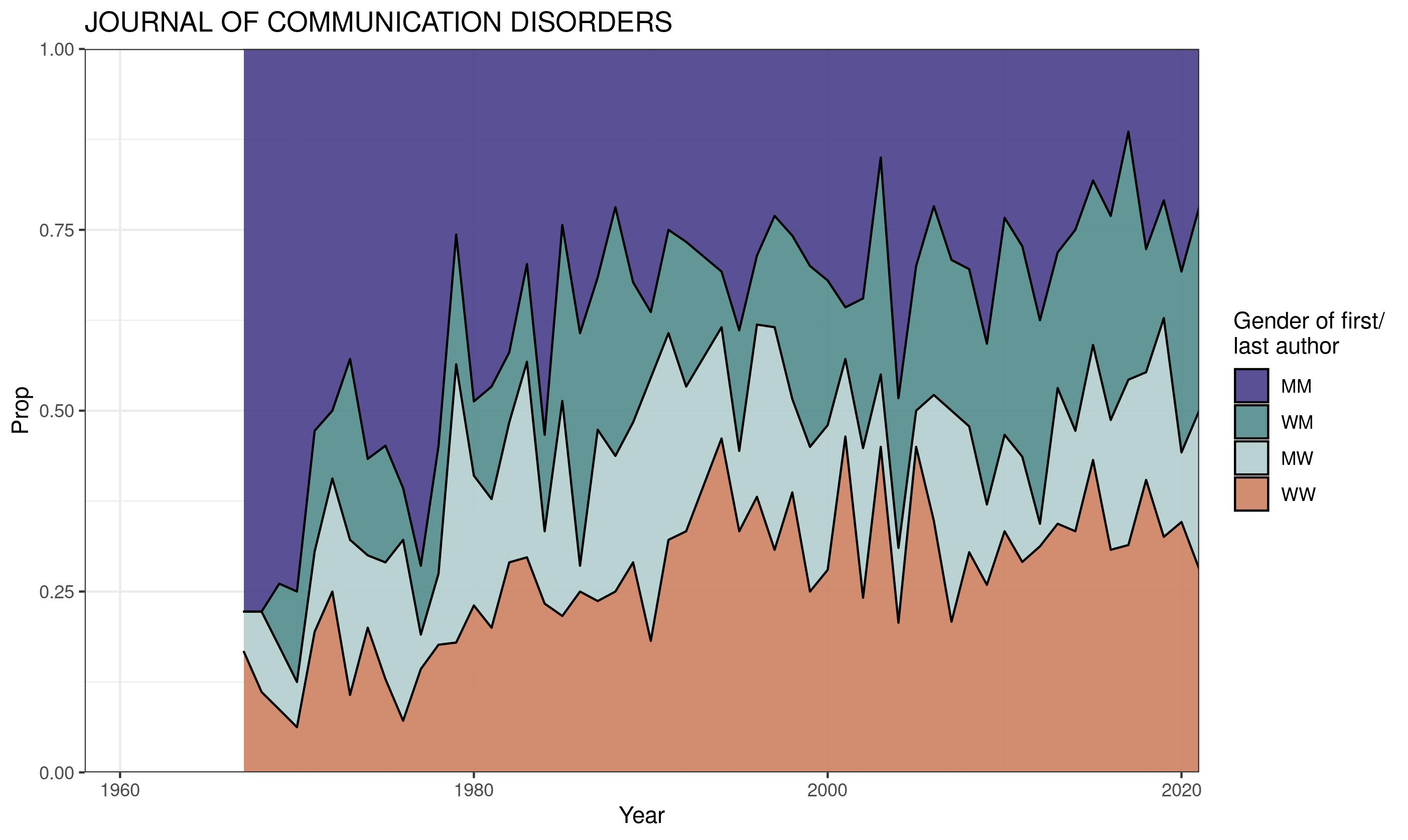

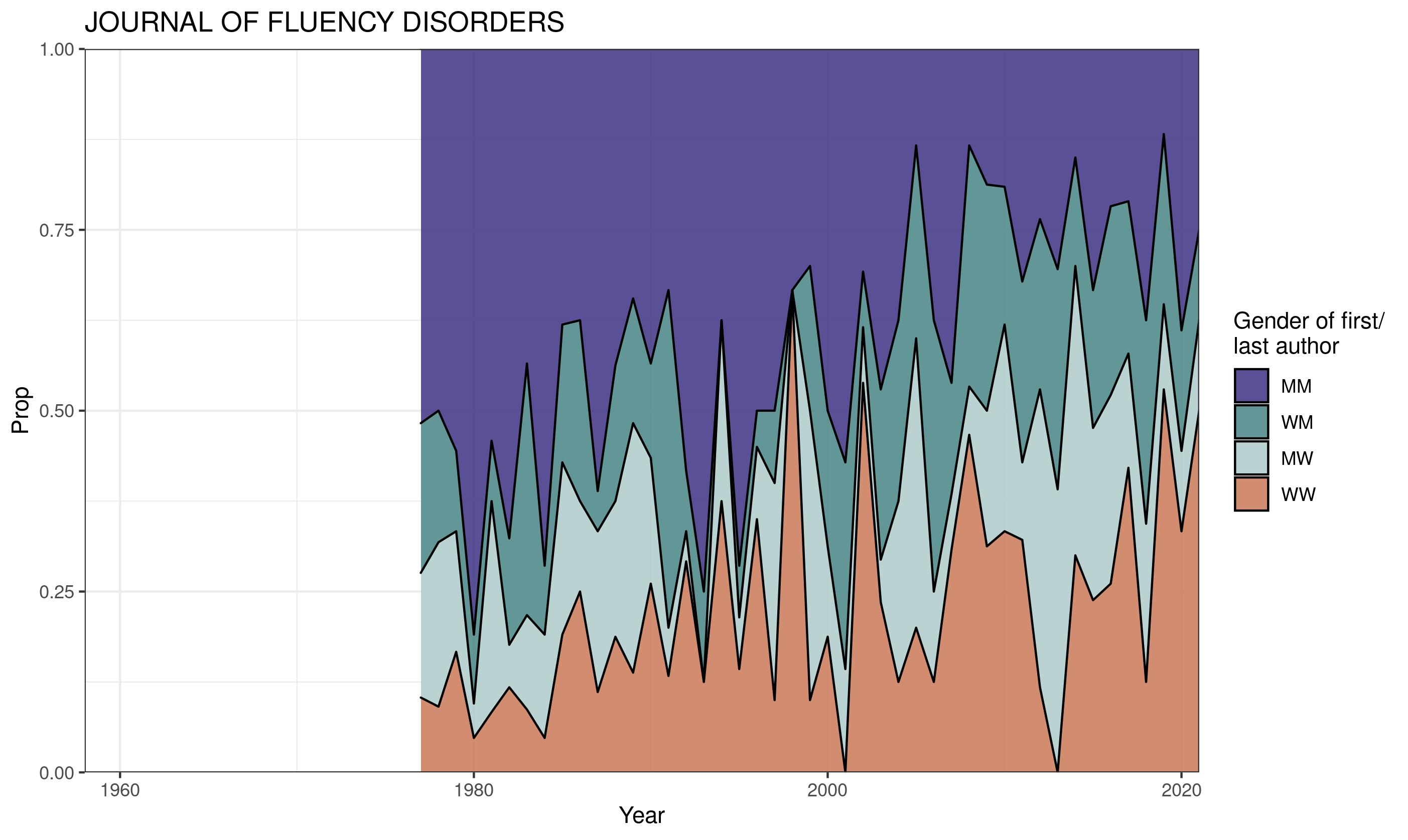

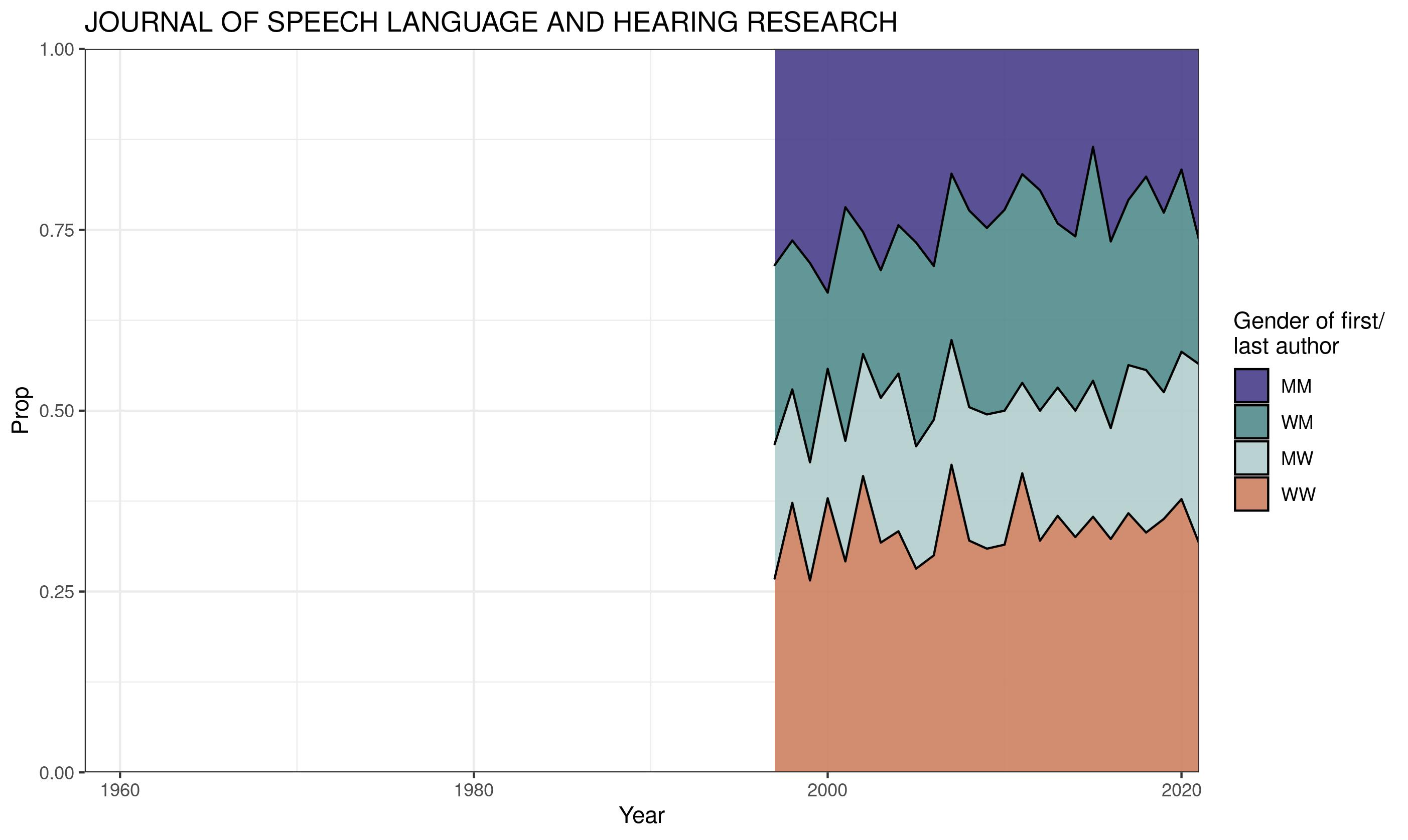

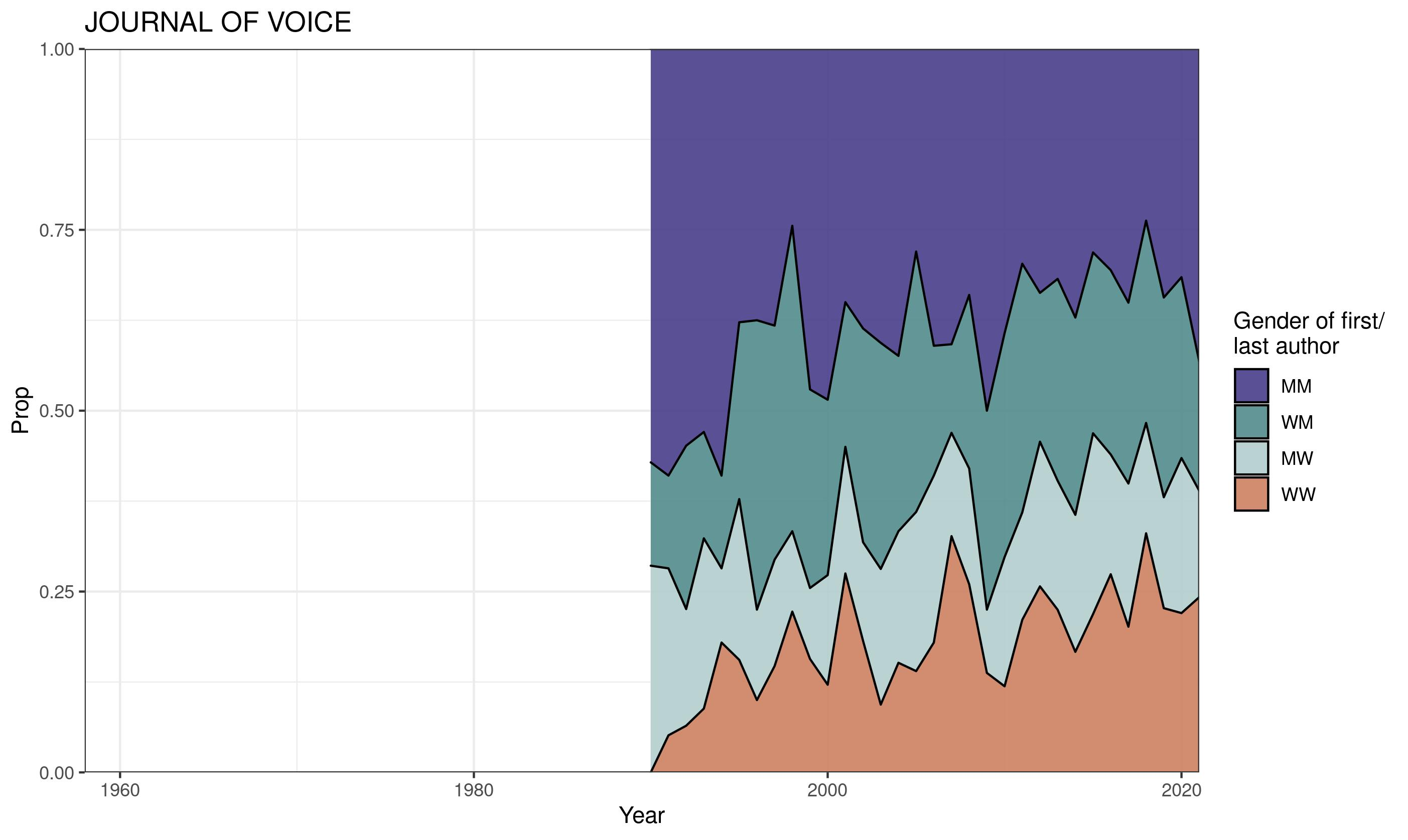

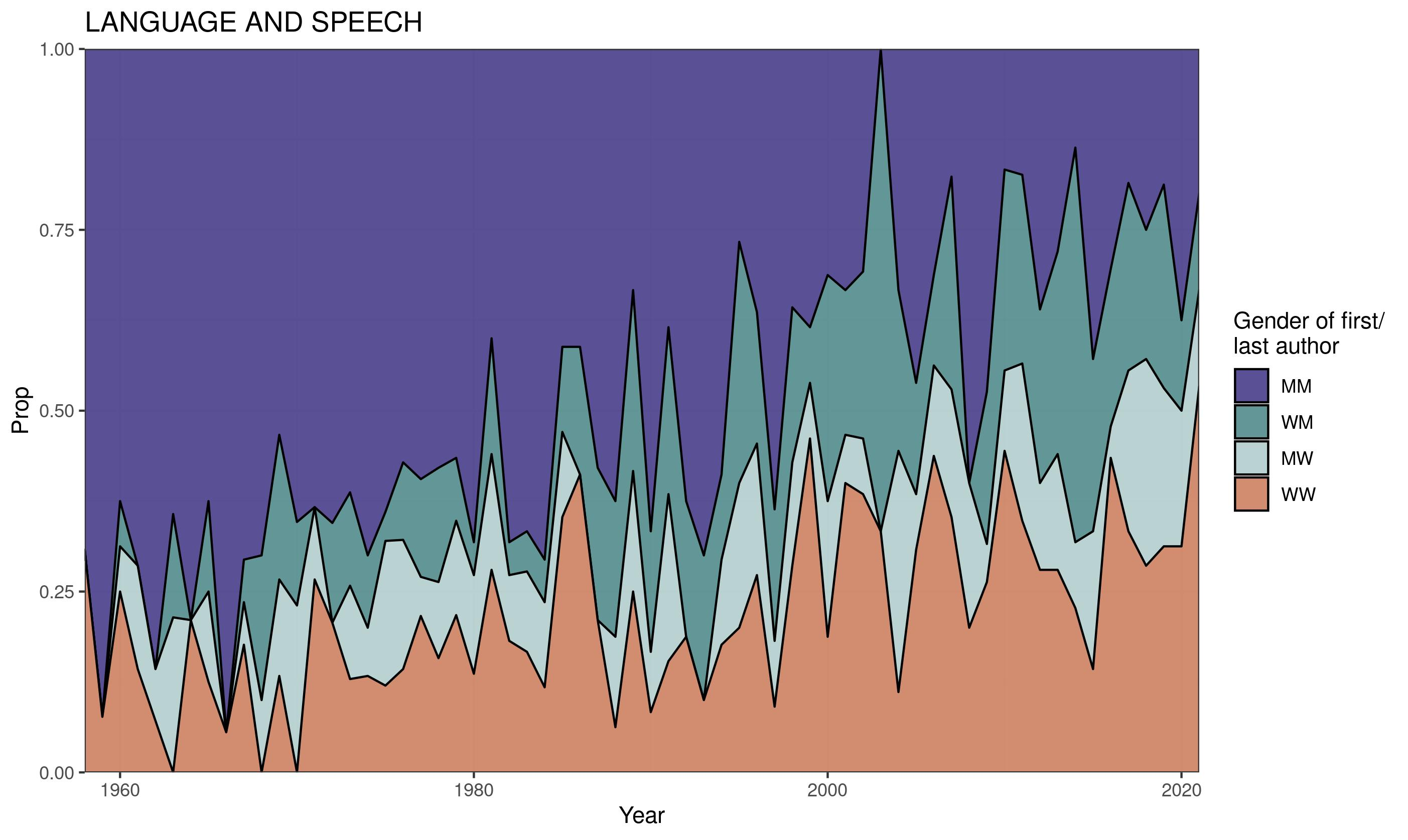

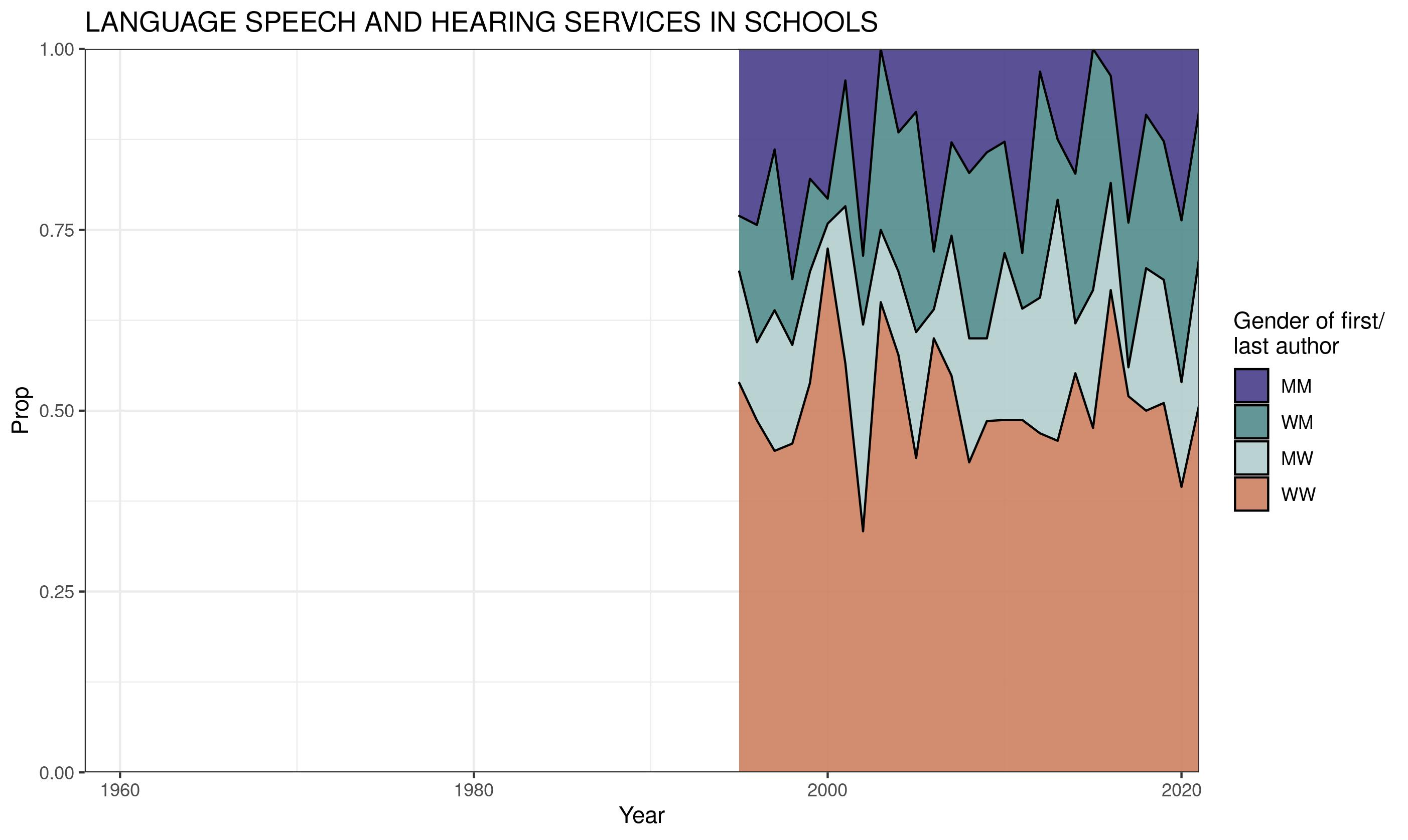

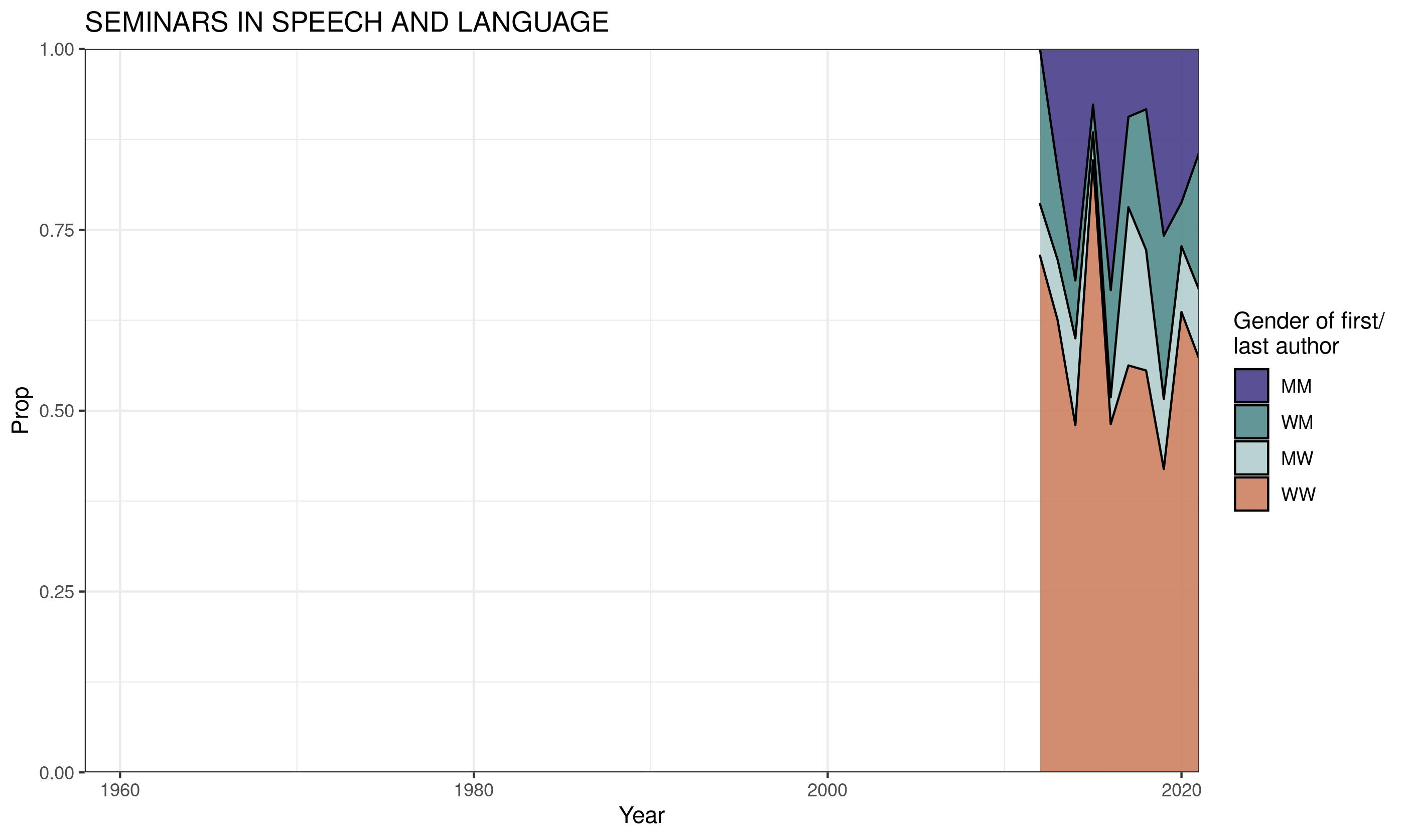

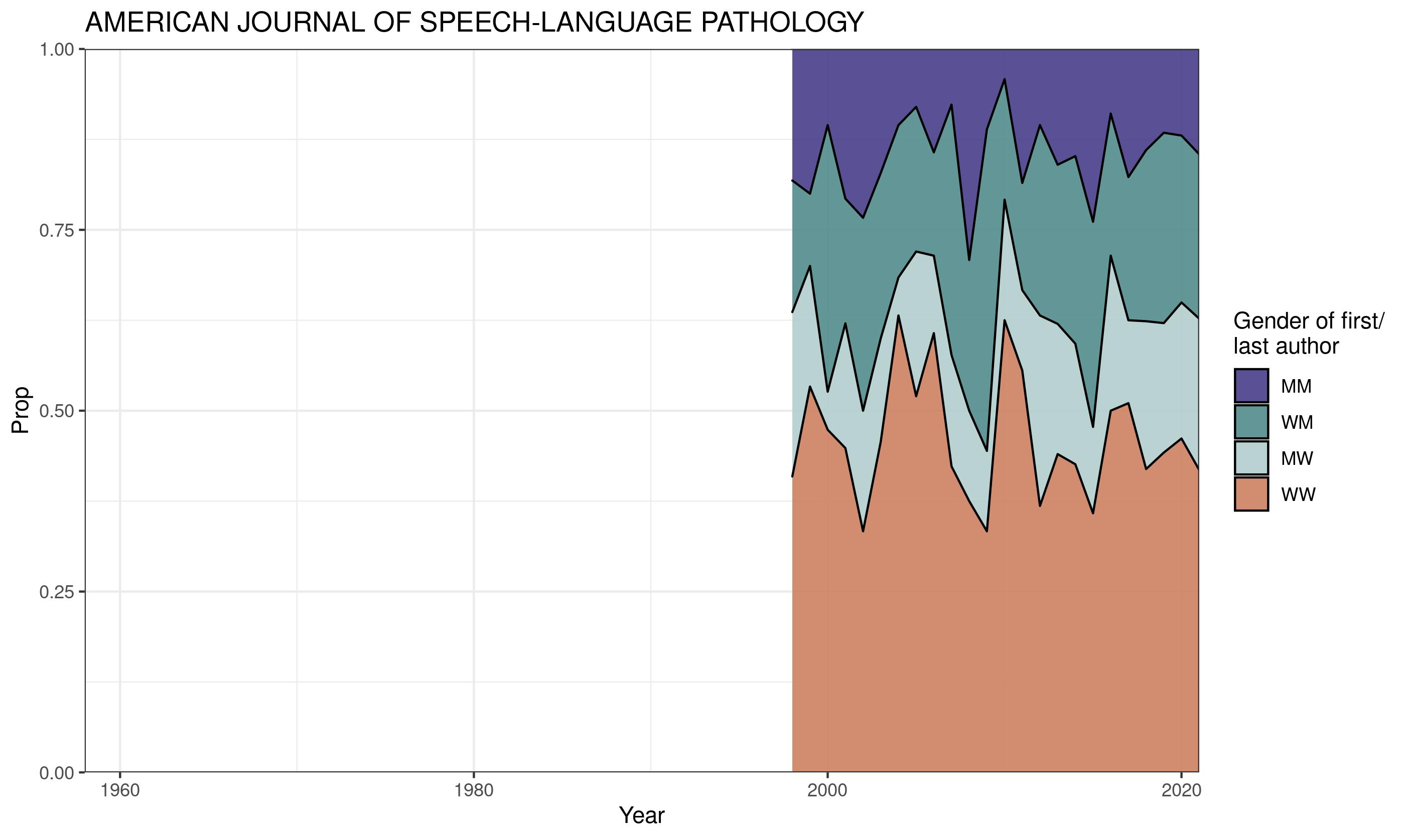
